# Differential Behavior of Conformational Dynamics in Active and Inactive States of Cannabinoid Receptor 1

**DOI:** 10.1101/2024.04.17.589939

**Authors:** Ugochi H. Isu, Adithya Polasa, Mahmoud Moradi

## Abstract

Cannabinoid receptor 1 (CB1) is a G protein-coupled receptor (GPCR) that regulates critical physiological processes including pain, appetite, and cognition. Understanding the conformational dynamics of CB1 associated with transitions between inactive and active signaling states is imperative for developing targeted modulators. Using microsecond-level all-atom molecular dynamics (MD) simulations, we identified marked differences in the conformational ensembles of inactive and active CB1 in *apo*. The inactive state exhibited substantially increased structural heterogeneity and plasticity compared to the more rigidified active state in the absence of stabilizing ligands. Transmembrane helices TM3 and TM7 were identified as distinguishing factors modulating the state-dependent dynamics. TM7 displayed amplified fluctuations selectively in the inactive state simulations attributed to disruption of conserved electrostatic contacts anchoring it to surrounding helices in the active state. Additionally, we identified significant reorganizations in key salt bridge and hydrogen bond networks contributing to the CB1 activation/inactivation. For instance, D213-Y224 hydrogen bond and D184-K192 salt bridge showed marked rearrangements between the states. Collectively, these findings reveal the specialized role of TM7 in directing state-dependent CB1 dynamics through electrostatic switch mechanisms. By elucidating the intrinsic enhanced flexibility of inactive CB1, this study provides valuable insights into the conformational landscape enabling functional transitions. Our perspective advances understanding of CB1 activation mechanisms and offers opportunities for structure-based drug discovery targeting the state-specific conformational dynamics of this receptor.

**Graphic for manuscript:** For Table of Contents Only

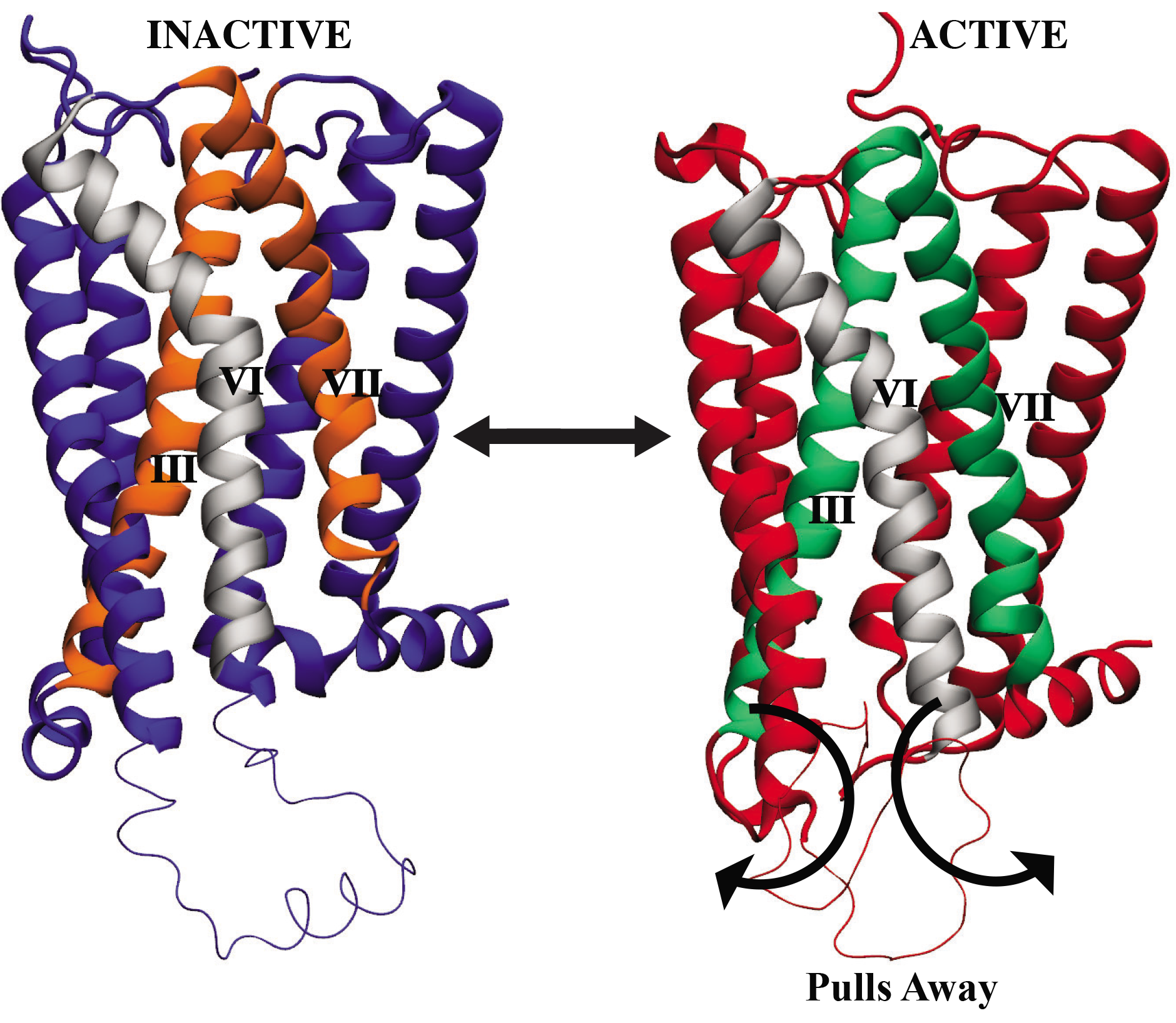

## Introduction

GPCRs are highly dynamic proteins^1^ that explore a broad spectrum of structural conformations, encompassing both their active and inactive functional states. Investigating the diversity in conformation and state-dependent dynamics is essential to understand how GPCRs are activated.^2–5^ Understanding activation mechanisms in GPCRs is a fundamental aspect of designing drugs based on GPCR structures.^6^ This is especially significant because over 30% of drugs available in the market are designed to target GPCRs.^7–10^ Crystal structures have offered valuable insights into the structure of CB1; however, they provide only a limited understanding of its activation mechanism, as they only capture static snapshots of the receptor. Complimentary experimental and computational techniques such as nuclear magnetic resonance (NMR) spectroscopy,^11,12^ hydrogen-deuterium exchange mass spectrometry (HDX-MS), and MD simulations,^13–17^ provide a dynamic views of receptor flexibility and the structural ensemble.^18–20^ To date, MD simulations have been instrumental in characterizing GPCR dynamics^21–23^ and enhancing interpretations of crystallographic data.^24,25^ All-atom unbiased MD simulations spanning microsecond to millisecond timescales can comprehensively sample conformational transitions.^9,26–28^ Numerous MD studies on GPCRs such as *β*2- adrenergic receptor and A2A receptor have identified ligand binding sites, activation mechanisms, and allosteric modulation.^26,29–31^ Specifically, comparing MD simulations in both the active and inactive states have highlighted conserved intramolecular interactions that stabilize GPCR conformations.^3,32–36^ Comparative MD studies of multiple functional states have provided insights into the conformational dynamics underlying CB1 function.^4,5,37–39^ Recent studies have shown that some GPCRs exhibit increased flexibility in the inactive state compared to the active state in apo form, contrary to the traditional understanding.^4,39–41^ For example, microsecond-timescale MD simulations of the *β*2-adrenergic receptor revealed greater conformational heterogeneity in the inactive state, while displaying active-like conformational elements.^42^

CB1 is a Class A GPCR that binds endogenous cannabinoids as well as exogenous ligands such as tetrahydrocannabinol (THC),^43^ the primary psychoactive constituent of cannabis.^44^ CB1 is one of the most abundant GPCRs in the central nervous system^45^ and a promising therapeutic target for pain, inflammation, mood control,^46^ obesity, neurodegeneration, and substance abuse disorders.^47–49^ However, chronic activation of CB1 is also associated with risks like addiction and psychosis. GPCRs such as CB1 contain 7 transmembrane (TM) alpha helices and signal via conformational changes between inactive and active states. In the inactive state, CB1 is stabilized by interactions between the TM3 (R214) and TM6 (D338) residues called the “ionic lock”.^37^ Upon ligand binding, this network is disrupted, enabling rearrangement of the helices to create a ligand binding pocket and expose residues for G protein coupling. Characterizing the conformational dynamics of CB1 in its inactive and active states is essential to understand its activation mechanisms as a GPCR and enable structure-based drug design efforts targeting this therapeutically important receptor. Crystal structures have been solved for CB1 in inactive and active states, providing snapshots of CB1 activation.^5,32^ To elucidate the intrinsic conformational dynamics of CB1, we performed multi-microsecond all-atom MD simulations of CB1 in apo inactive state and apo active state. Two independent 1 *μ*s simulations were run for each state in explicit membrane/solvent environment, totaling 4 *μ*s of aggregate sampling.

GPCRs such as CB1 rely on conformational flexibility to transduce signals regulating physiology. However, details on dynamics differentiating inactive and active states require further elucidation to inform targeting. Our molecular dynamics simulations revealed greater conformational heterogeneity in the inactive state compared to the active state. These findings are consistent with the observations made by Sangho Ji et al.,^4^ which also noted increased flexibility in the inactive state of CB1 when toggle switches were mutated.

Notably, our simulations identified the significant role of TM7 in driving the structural differences between active and inactive states of CB1, forming stabilizing interactions specifically in the active state. Previous research has identified a set of switches within TM7 residues that facilitate the transmission of signals from the ligand binding site to the G protein coupling region.^50^ During the activation process of CB1 and most GPCRs, a critical event involves the inward movement of TM7.^3,51^ This movement is triggered when the ionic lock, a structural element within the receptor, is disrupted, causing the TM3 and TM6 helices to move apart.^50^ Our findings provide atomistic details into the functional role of TM7 in the activation mechanism of CB1. Additionally, our comparative analysis highlighted the roles of electrostatic interactions in the distinct conformational dynamics between states. The inactive state contains a dense network of electrostatic interactions that rigidify the conformation of ECL1, the N terminus, and helices 1, 2 and 6 TM bundles.^5^ This network is formed by an array of polar contacts and charge-charge interactions between conserved motifs and microswitches associated with CB1 function. This is disrupted in the active state, enabling TM movements necessary for activation.^52,53^ Therefore, these electrostatic interactions could constitute an integral regulatory mechanism controlling state-dependent CB1 dynamics. Our study sheds light on the dynamic context of the suggested rotating network of extracellular electrostatic interactions influencing CB1 function.

In summary, our comparative MD simulations enhance our understanding of CB1 activation mechanisms. We observe that the inactive state displays greater conformational plasticity compared to the more constrained active state, with TM3 and TM7 playing significant roles in this regard through extensive coupling interactions. Our study progresses our understanding of the conformational landscape governing CB1 dynamics. Our atomistic insights could guide interpretation of new CB1 structures and design of functionally selective drugs targeting this important receptor.

## Methods

### Molecular Modeling and Simulation Systems

We employed all-atom MD simulations to elucidate the conformational dynamics of the CB1 receptor, within a modeled membrane environment. We designed two CB1 simulation systems based on the high-resolution crystal structures of human CB1 representing the active state (PDB entry: 5XRA, 2.80 Å resolution)^37^ and the inactive state (PDB entry: 5TGZ, 2.95 Å resolution).^5^ To ensure accuracy of the simulations, four residues in the crystal structures (A210T, K273E, V283T, and E340R) (Fig. 1) were mutated back to the wild-type amino acid residues. For both systems, we utilized a Monte Carlo algorithm to model the missing loop regions using the program Modeller,^54^ with active crystal structures modeled between residues 307-336, and inactive crystal structures modeled between residues 307-331 (Fig. 1) connecting TM5 - TM6. The prepared CB1 proteins were embedded in a 90% hydrated 1-palmitoyl-2-oleoyl-glycero-3-phosphocholine (POPC) and 10% cholesterol lipid bilayer to mimic the native membrane environment using the Membrane Builder module in CHARMM-GUI,^55^ and 0.15 M NaCl included (in addition to the counterions used to neutralize the protein) to mimic physiological conditions. The simulation systems was solvated in a cubic box of TIP3P water, with dimensions of 150 Å *×* 150 Å *×* 150 Å. In the first and second simulation runs, the total number of atoms in the inactive state systems were 70,985 and 71,255, respectively. Meanwhile, in the active state systems, there were 78,565 and 79,255 atoms for the first and second simulation runs. All MD simulations were performed in NAMD 2.13^56^ using the CHARMM36m force field^57^ for proteins, lipids, and ions. Energy minimization was initially applied to both the active and inactive state systems using the conjugate gradient algorithm for 10,000 steps. Production runs were then carried out for 1 *μ*s for 4 systems, totalling 4 *μ*s. The equilibration phase was performed in an NVT ensemble, while the production runs were conducted in an NPT ensemble. A time step of 2 fs was utilized, and the temperature was maintained at 310 K using a Langevin thermostat. The Nośe-Hoover Langevin piston method^58^ was used to control pressure at 1 bar.

**Fig. 1.**
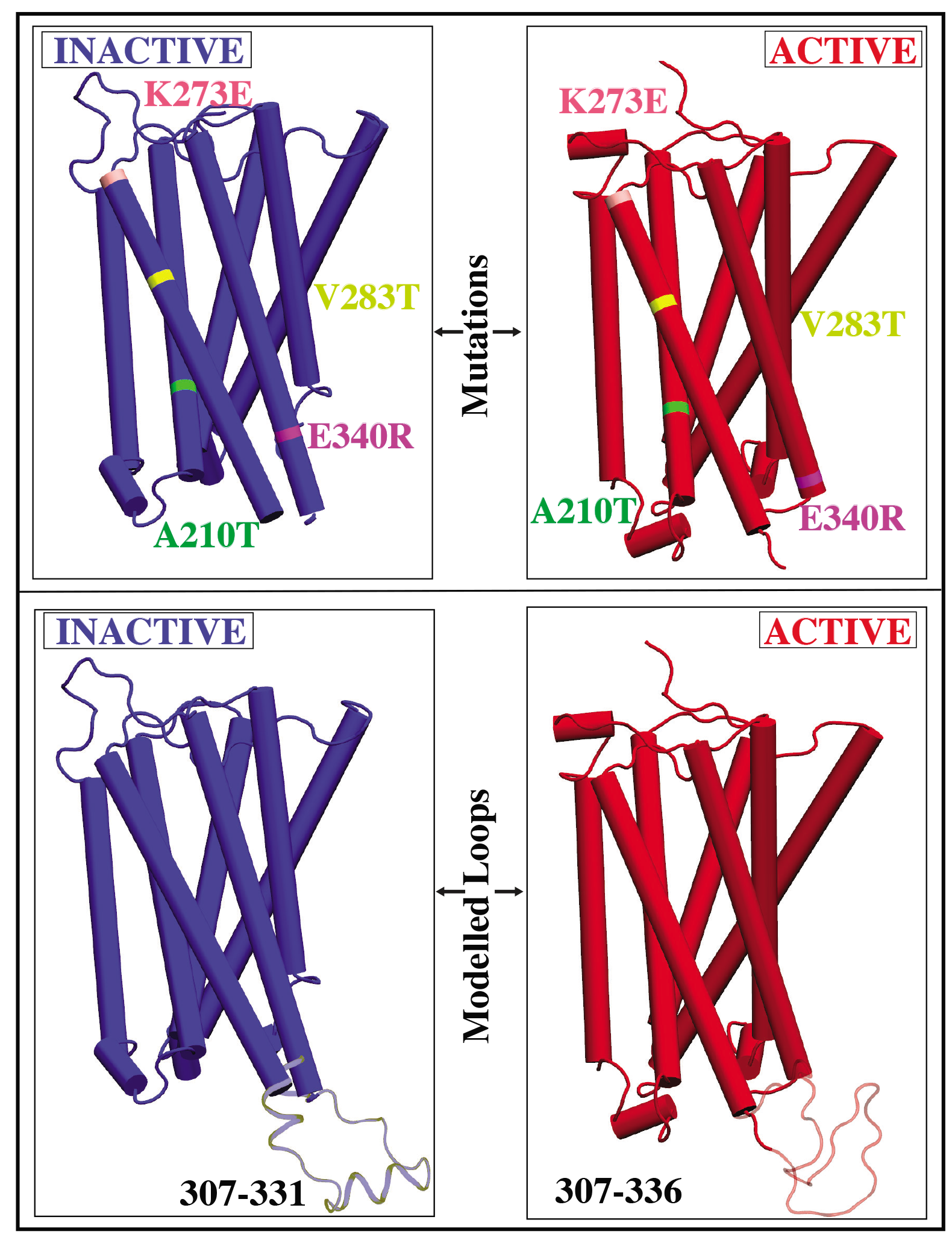
Introduced mutations and loop modeling in CB1 receptor for active and inactive state. E340R, A210T, V283T, and K273E, reverted to their original wildtype sequences and missing extracellular loop 2 segments modeled.

### Trajectory Analysis

The TM helices and other subdomains were defined as follows using published methods^4,59^ : N-terminal region (residues 104 - 112), TM1 (residues 113 - 148), ICL1 (residues 149 - 152), TM2 (residues 153 - 180), ECL1 (residues 181 - 185), TM3 (residues 186 - 219), ICL2 (residues 220 - 229), TM4 (residues 230 - 253), ECL2 (residues 254 - 273), TM5 (residues 274 - 306), ICL3 (residues 307 - 336), TM6 (residues 337 - 369), ECL3 (residues 370 - 372), TM7 (residues 373 - 401), TM8 (residues 402 - 412), and C-terminal region (residues 413 - 414). The root-mean-square deviation (RMSD) trajectory tool in VMD^60^ was employed to calculate the RMSD, with C*α* atoms considered for these calculations. The average RMSD was determined based on the entire trajectory, and error bars were used to indicate the standard deviation in the data. To assess the flexibility of individual residues, the root-mean-square fluctuation (RMSF) was calculated using C*α* atoms, aligning the trajectory against the crystal structure. Salt bridge interactions were identified using the VMD timeline plugin,^60^ with a cutoff distance of 4 Å. The VMD salt bridge plugin enabled the calculation of the distance between two charged residues throughout the simulation, specifically measuring the distance between the oxygen atom of the acidic residue and the nitrogen atom of the basic residue. Principal component analysis (PCA) was performed using the PRODY software,^61^ considering only C*α* atoms for the calculations. Hydrogen bond (H-bond) analysis was conducted using the VMD HBond plugin,^60^ with a cutoff distance of 3.5 Å and an angle cutoff of 30°.

## Results and Discussion

### Protein Dynamics: Exploring Conformational Variability in Different States

Our comparative MD simulations demonstrated significant differences in the behavior of the apo CB1 receptor between its active and inactive states. By employing RMSD and RMSF calculations, we were able to analyze the structural dynamics of CB1 in both states, revealing that the inactive state of CB1 seems to adopt a wider variety of conformations compared to its active state in the absence of ligand interactions.

RMSD values of the TM domain remained below 6Å throughout the active state simulations, indicating minimal deviation from the initial crystal structure after agonist dissociation (Fig. 2A). The consistent RMSD profile reflects sustained rigidity and structural stability of the active state despite the loss of agonist interactions, likely due to inherent intramolecular interactions between the TM helices maintaining an active-like ensemble. In contrast, the RMSD of the inactive state steadily increased above 9Å with larger fluctuations, suggesting continuous exploration of diverse conformations diverging significantly from the initial crystal structure (Fig. 2B). This deviation from traditional models where inactive GPCRs maintain restrained ensembles stabilized by conserved interactions indicates that the ensemble of inactive CB1 structures differs from the antagonist-bound crystal conformation. The heightened RMSD in the inactive state suggests increased flexibility necessary to sample intermediates competent for G protein coupling after antagonist release, potentially representing transitional intermediate structures explored during the activation process. Evaluation of per-residue fluctuations revealed differences in dynamics between the active and inactive states. RMSF profiles showed enhanced flexibility in extracellular and intracellular loop regions in both states compared to the rigid TM cores (Fig. 2C-D). However, distinct dynamics were observed in specific structural elements. In the active state, the TM4-5 interhelical loop exhibited greater rigidity compared to the inactive state, while TM7 showed slightly higher flexibility in the inactive state (Fig. 2C-D).

**Fig. 2.**
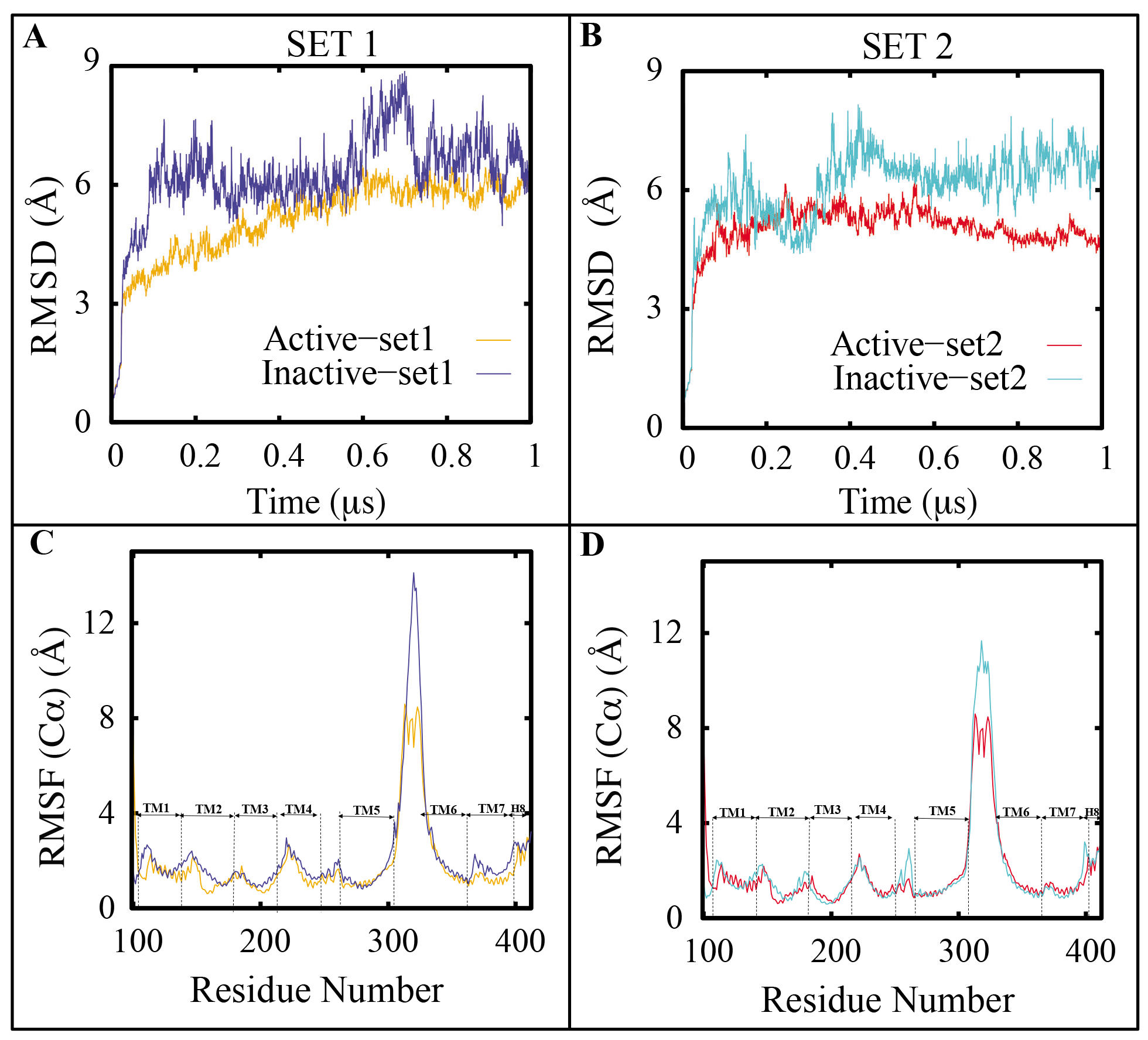
Analysis of the CB1 structural stability in the active and inactive states. (A-B) RMSD of the CB1 receptor in the active and inactive states from two independent sets of simulations. Similarly, RMSF of each residue is shown in (C-D) for the same simulations.

To further probe the conformational dynamics, we employed PCA.^61^ This approach played a crucial role in streamlining our dataset, effectively capturing the intricate trajectory details while reducing dimensionality. Projection of the trajectory data onto the first two principal components showed clear separation between inactive and active states (Fig. 3). Strikingly, PCA revealed the inactive state consistently displayed substantially greater motion along the first principal component, even when comparing independent simulation sets (Fig. 3A-C). This suggests that the inactive state ensemble explores a wider conformational space through collective structural rearrangements.

**Fig. 3.**
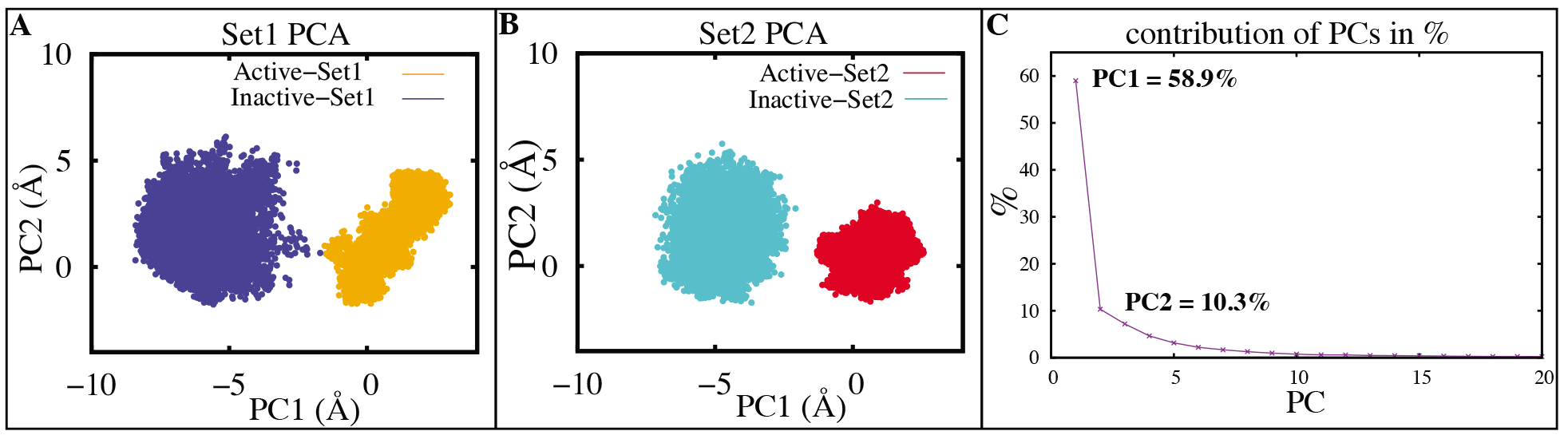
Conformational regions associated with the active and inactive states of CB1, as shown by PCA. (A,B) Projections of PC1 and PC2 for models of CB1 inactive (blue, cyan) and active (orange, red) states, for both replicates (C) The top 20 PCs’ percentage of variance.

The pronounced increased flexibility of the inactive state could potentially have significant functional implications. GPCRs such as CB1 undergo concerted conformational changes upon ligand binding to initialize downstream signaling cascades. Our observation of enhanced inactive state dynamics likely reflects an intrinsic predisposition to readily adopt various conformations in response to ligand binding. This enables transitions between signaling states, a process that appears inherently more hindered from the rigid active state. The complex conformational landscape explored by inactive CB1 suggests a degree of conformational selection upon ligand binding. Certain conformational intermediates within the broad inactive ensemble may be preferentially stabilized and selected by different ligands. This enables modulation of downstream signaling through differential ensemble stabilization. Our findings indicate inactive CB1 may utilize conformational selection, although this mechanism may be combined with the induced fit mechanism.

Therefore, our MD simulations could advance our understanding of the conformational dynamics distinguishing CB1 states. We observe that contrary to traditional models, the inactive CB1 explores a wider conformational landscape and exhibits greater flexibility in apo conditions, compared to the active state. The intrinsic malleability of inactive CB1 likely primes it for conformational transitions during activation.

### TM7’s Pivotal Role in Conformational Differences Between Active and Inactive CB1

To elucidate the specific structural elements contributing to these differences, we conducted detailed analyses of individual TM helix motions. These investigations highlighted the significant roles of TM3 and TM7 in modulating state-dependent CB1 dynamics. Analysis of TM helix RMSD uncovered notable conformational changes in TM3 and TM7 between states. In the first simulation set, TM3 exhibited higher flexibility in the active state, with RMSD climbing to 2Å compared to 1.2Å when inactive (Figs. 4 and S1). This confirms that TM3 undergoes substantial structural rearrangements upon activation.^3,4^ NMR studies of the *β*2 adrenergic receptor corroborate this, showing TM3 displayed distinct chemical shift perturbations between inactive and active states.^62^ Interestingly, this particular conformational change of TM3 was exclusive to the first set of our simulations, and not observed in the second set (Fig. S1).

**Fig. 4.**
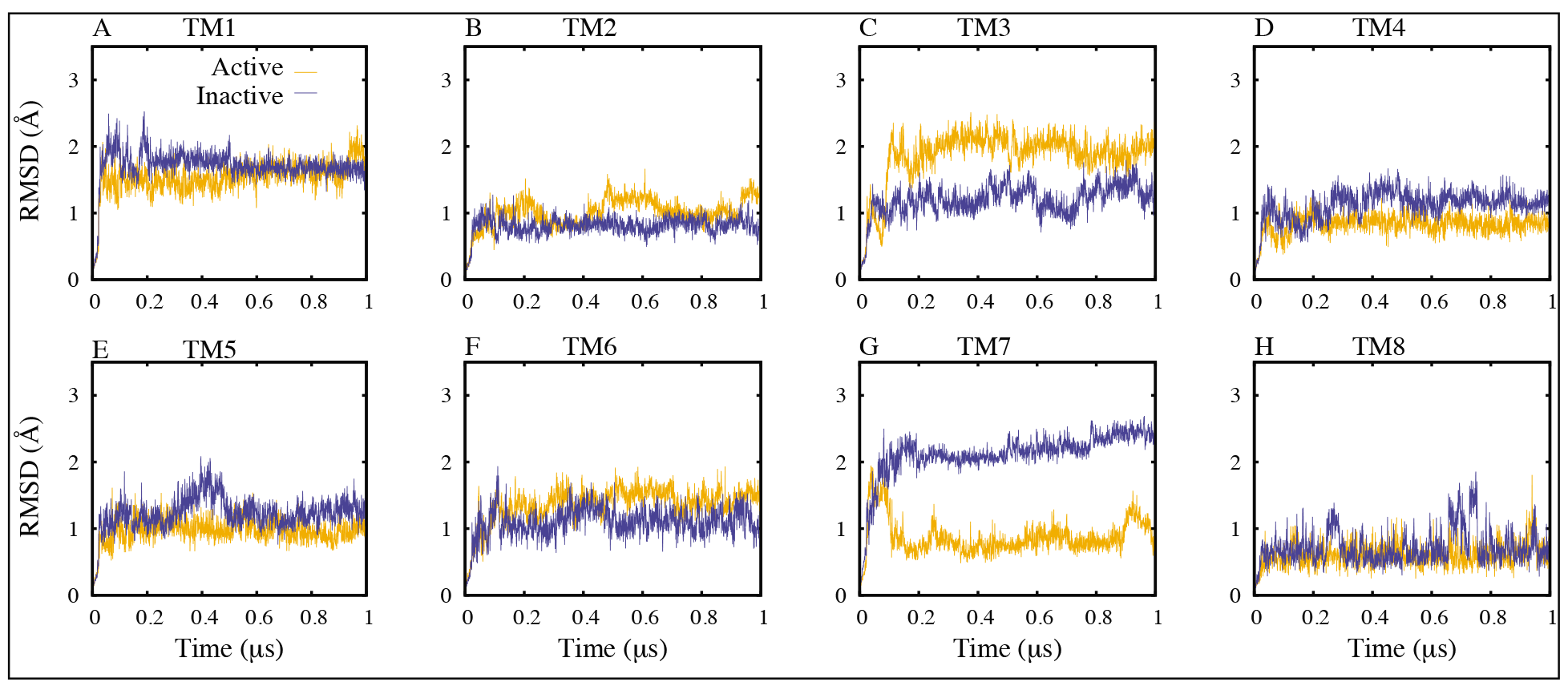
RMSD Profiles of TM Helices in Set 1. Highlighting conformational changes in TM3 and TM7 between active (orange) and inactive (blue) states. The figure depicts RMSD analyses of the TM helical bundles for set 1 simulation.

In contrast, TM7 displayed enhanced flexibility in the inactive state in both independent simulations, with RMSD elevated compared to the active state (Figs. 4 and 5A). The pronounced RMSD fluctuations signifies the substantial conformational malleability of TM7 specifically when CB1 adopts the inactive state. TM7 forms extensive stabilizing intramolecular contacts with other helices when activated to enable its inward movement.^3,63–65^ Thus, our findings agrees with biochemical studies showing TM7 rearrangements are integral to the activation mechanism of Family A GPCRs such as CB1.^39,66–69^ To further examine these trends, we calculated the interhelical angles formed between TM helices as described in Methods. This confirmed the increase in angle fluctuations for inactive state TM7 across both simulations (Fig. 5B). TM1-TM7 angle variations corroborated the enhanced flexibility of inactive state TM7. Quantifying angle changes provides additional evidence that inactive undergo marked structural rearrangements to facilitate CB1 state transitions.

**Fig. 5.**
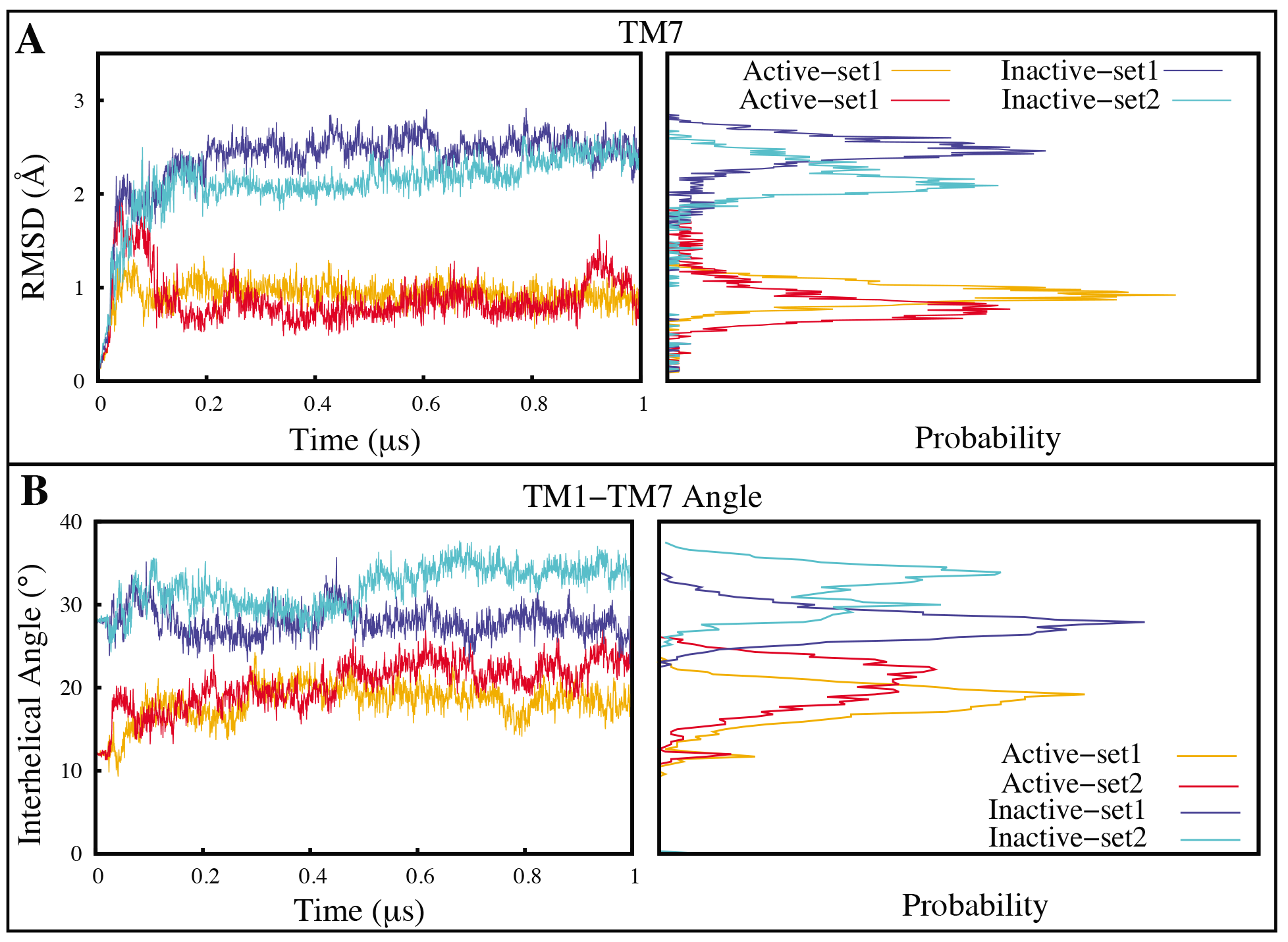
Structural variance between the CB1 active and inactive states is significantly in- fluenced by local conformational changes. (A) The RMSD of the TM7 helix in both active (orange and red) and inactive states (blue and cyan) (B) Interhelical angle between TM1 and TM7 in the active (orange and red) and inactive (blue and cyan) states. All graphs also display the probability density distribution.

The functional significance of TM3 and TM7 dynamics arises from their known involvement in GPCR activation processes.^3,39,51^ Agonist binding disrupts the ionic lock between TM3 and TM6, enabling TM movements required for G protein coupling.^69–71^ Our findings indicate that the increased flexibility of TM3 in the active state facilitates its outward movement, resulting in the disruption of the ionic lock. Moreover, our observations reveal that TM7 forms a H-bond with TM2, suggesting that it potentially stabilizes its inward movement in the active state. The observed dynamics of TM7 are likely to facilitate rearrangements that disrupt the inactive state interactions, ultimately leading to an intermediate conformation resembling the active state. TM3 and TM7 seem to have distinct functions in each activation state, enabling the transition of CB1 conformations. TM3 maintains stability in the inactive state via the ionic lock, while it gains flexibility during activation to disrupt the ionic lock. Meanwhile, TM7 becomes rigid post-activation, following the loss of its coupling interactions. Our study delves into the detailed interplay between TM3 and TM7, shedding light on their roles in state transitions at an atomic level. Another significant observation from our analysis is that TM7 rearrangements are integral not only for activation, but also deactivation processes. The marked flexibility of inactive TM7 likely enables the precise conformational changes necessary to impede further signaling and stabilize the inactive state. Our analysis validates that TM7 rearrangements are critical in both directions between functional states. This aligns with NMR investigation and biochemical evidence, affirming that TM7 serves as a central element influencing the conformational adaptability among CB1 models.^72^

Elaborating on the distinct role of TM7 within the dynamic behavior of CB1, further analysis uncovered a key H-bond interaction between Ser 390 in TM7 and Asp 163 in TM2 that appears crucial for stabilizing the active state. Asp 163 is known to coordinate sodium binding, highlighting interplay between the ionic and H-bond networks regulating CB1 activation.^4,73^ The Ser 390 - Asp 163 H-bond exhibited contrasting behavior between states across both simulations. In the active state, the H-bond formed early on and remained stable throughout the multi-microsecond timescale (Fig. 6A,C). The persistence of this interaction implies it constitutes a stabilizing factor maintaining the active conformation. This agrees with biochemical studies showing TM7 rearrangements are integral to constraining the active state.^66–69^ However, for the inactive state, this H-bond showed instability and partial dissociation (Fig. 6B,D and Mov. S1). The shift in H-bond stability underscores its role as a sensor of CB1’s conformational state. Notably, previous computational and crystallographic studies have identified this structurally conserved H-bond, but lacked dynamical context into its state-dependent behavior.^4^ Our simulations offer insights into how the Ser 390 - Asp 163 interaction switches between stabilized and destabilized states to regulate CB1 activation processes. We propose that the disruption of this H-bond is an early event that enables TM7 outward movement to facilitate transition to the inactive state. Our observation of this interaction’s instability in the inactive state agrees with the model of GPCRs undergoing substantial conformational changes upon activation state transitions.^5,37,63–65^ This likely arises from reorganization of the H-bond network, enabling rearrangements like the rotamer toggle switch of Ser 390. The contrasting dynamics of the Ser 390 - Asp 163 H-bond aligns with this understanding of key interactions acting as switches to control state transitions.^63–65^ Therefore, our study sheds light on a highly conserved H-bond interaction exhibiting specialized state-dependent dynamics that appears vital for modulating CB1 activation. The stability of this TM7-mediated contact in the active state relative to its marked instability when inactive reveals its role as a conformational switch controlling CB1 function. Our findings offer atomistic insights into how dynamic rearrangements in H-bond networks enable functional transitions of CB1, advancing our understanding of the intricate molecular mechanisms involved in CB1 activation processes and signaling.

**Fig. 6.**
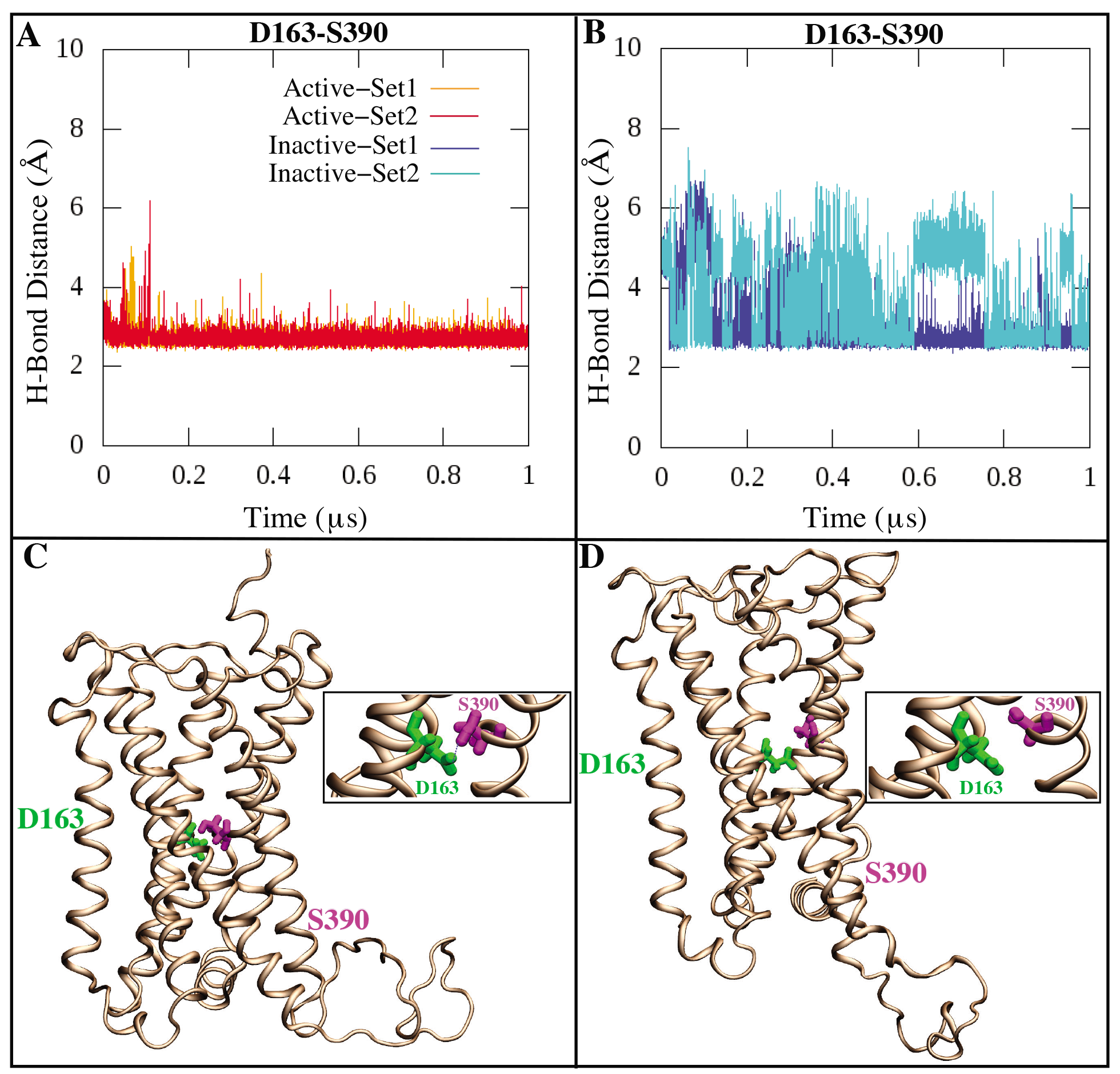
Hydrogen bond interaction between D163 and S390 stabilizes the TM7 helix in the active state of the CB1 receptor. (A-B) Time series of the hydrogen bond distances between the C*α* atoms of D163 (TM2) - S390 (TM7). (C-D) A graphic representation of hydrogen bond interactions between the D163 and S390 in the active and inactive states of CB1, respectively.

### Key Roles of Electrostatic Interactions in Conformational Dynamics Between States

Our microsecond-scale MD simulations revealed a dual salt bridge interaction network centered on Lys 192 in TM3 that displays distinct state-dependent dynamics between inactive and active CB1. Specifically, Lys 192 concurrently forms salt bridges with Asp 184 in extracellular loop 1 and Asp 176 at the extracellular end of TM2 when CB1 adopts the inactive state. This dual Asp 184 - Lys 192 and Asp 176 - Lys 192 salt bridge arrangement has been consistently observed in multiple crystal structures of inactive CB1. ^4,5,37,74,75^ Our simulations found that this paired salt bridge network was exceptionally stable over the multi-microsecond trajectory when CB1 maintained the inactive conformation in both independent simulation sets (Fig. 7B,E,F). In contrast, in the active state, the dual salt bridge arrangement was disrupted in our simulations (Fig. 7A,C,D and Mov. S2). The Asp 184 - Lys 192 salt bridge dissociated early in the trajectory within 200 ns in one simulation set, while it persisted throughout the multi-microsecond timescale in the other set (Fig. 7D). However, the Asp 176 - Lys 192 salt bridge showed distinct dynamics between the two independent active state simulations. It consistently ruptured within the first 200 ns in one set but remained weakly stable for most of the trajectory in the other set (Fig. 7A and Mov. S2). These variances highlight the increased flexibility of the dual salt bridge network in the active state ensemble. Notably, Lys 192 interconverted between pairing with Asp 176 and Asp 184, indicating rearrangement of the network. This suggests that the increased flexibility of Lys 192 enables the extracellular loop movements required to reach active-like conformations observed in our simulations.

**Fig. 7.**
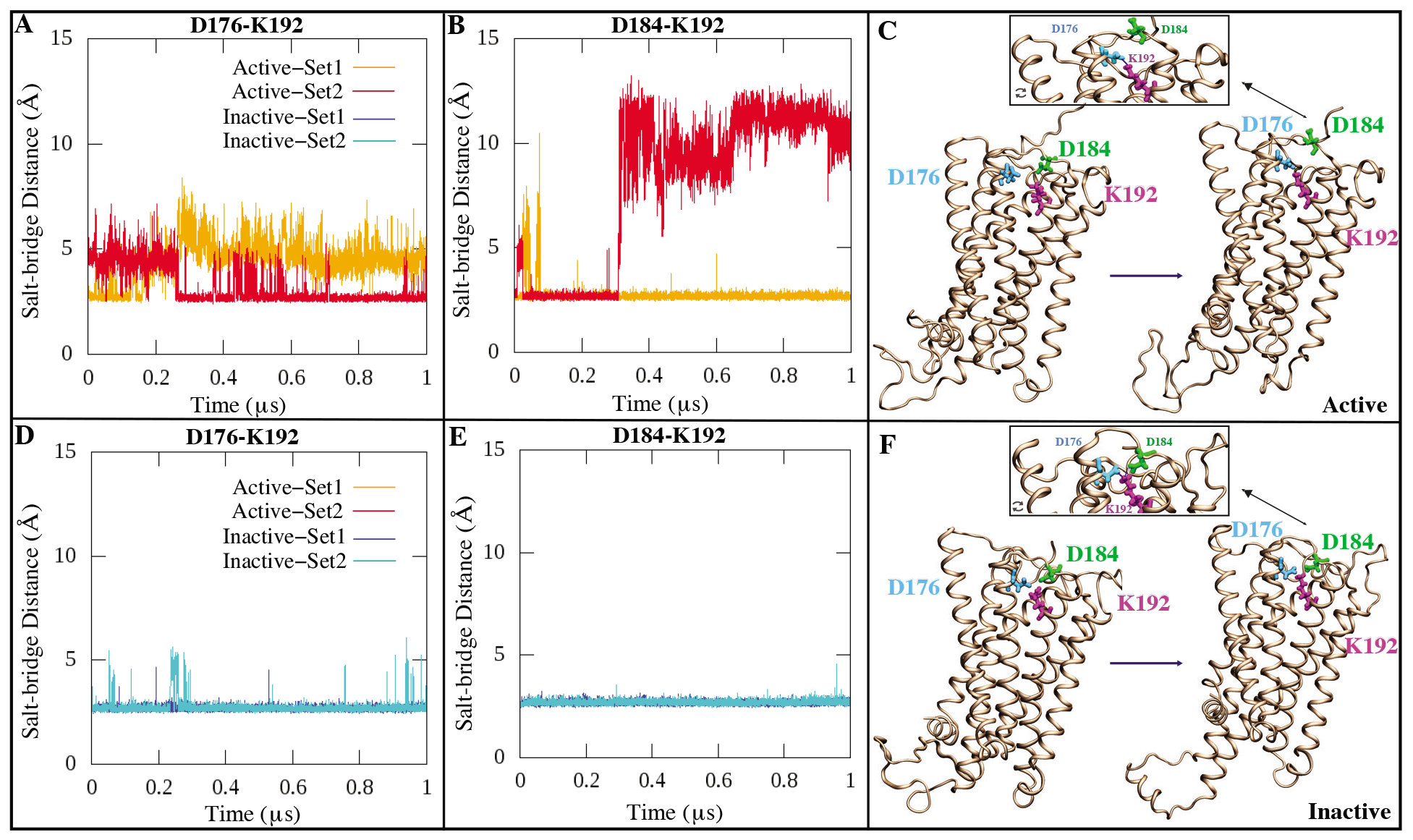
Time series of the salt bridge network between C*α* atoms of K192 (magenta), D176 (cyan), and D184 (green) in the (A, B) active and (D, E) inactive CB1 states across replicates. (C, F) Graphical illustration of K192-D176/D184 salt bridge interactions in the active and inactive CB1 conformations.

Aditionally, we identified a notable salt bridge formed specifically between Asp 213 in TM3 and Arg 230 in TM4 that was uniquely present in one active state simulation (Fig. 8D). This Asp 213 - Arg 230 salt bridge distinctly formed around 300 ns and remained stably intact throughout the multi-microsecond trajectory when CB1 was in the active state (Fig. 8D). It was completely absent in both inactive state simulations (Fig. 8E), indicating it is a specialized interaction associated with stabilization of activated CB1. Asp 213 is a crucial component of the well-preserved “ionic lock” network involving TM3, TM6, and TM7 that undergoes rearrangement upon activation, with the key ionic lock residues primarily consisting of Arg 214 and Asp 338.^5,37,73,76–78^ In the inactive state simulations, we observed that Asp 213 formed a persistent H-bond with Tyr 224 throughout both multi-microsecond timescales (Fig. 8B and Mov. S3). This conserved Asp 213 - Tyr 224 H-bond was present in the starting crystal structures of both active and inactive states. However, in one active state simulation, Asp 213 - Tyr 224 H-bond dissociated after 100ns to enable formation of the novel salt bridge with Arg 230, while the other active state simulation maintained the Asp 213 - Tyr 224 H-bond stably throughout (Fig. 8A and Mov. S3). The formation of the Asp 213 - Arg 230 salt bridge selectively when Asp 213 - Tyr 224 dissociated provides evidence it is a signature contact of the stabilized active CB1 state. Previous extensive research has suggested that Tyr 224 is a pivotal residue within the orthosteric binding site. It forms H-bonds with certain ligand residues and undergoes a reorientation of its sidechain, facilitating CB1 activation.^77,79,80^ Our observation of Asp 213 - Tyr 224 destabilization coupled to formation of Asp 213 - Arg 230 agrees with this model. Specifically, breaking of Asp 213 - Tyr 224 would allow Tyr 224 to sample alternative rotameric states to facilitate adoption of an active-like conformation, concurrently enabling the compensatory salt bridge with Arg 230. We also identified a unique salt bridge network involving Asp 104, Glu 106, and Arg 376 that changes significantly between different states (Fig. S2), alongside a H-bond between Ser 265 in ECL2 and Asp 366 in TM6 (Fig. S3). These molecular arrangements, not seen in the crystal structure, emerge in the inactive state simulations, maintaining presence predominantly in these states but disappearing in the active state of CB1. This analysis underscores the importance of considering the dynamic nature of CB1 and proteins at large, over relying solely on static structures, offering novel insights into the regulatory mechanisms of CB1 activation through conserved electrostatic interactions.

**Fig. 8.**
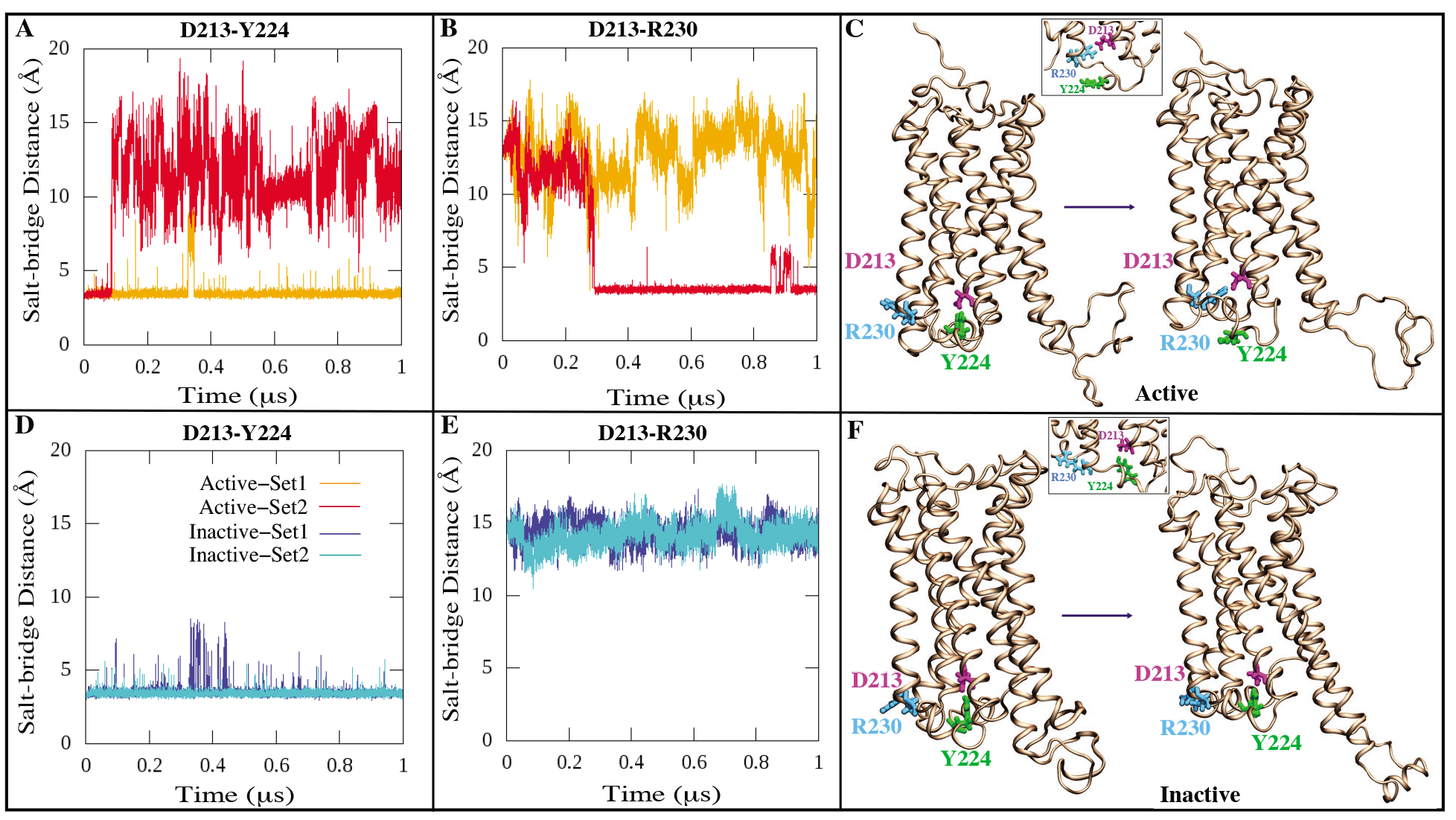
Network of salt bridge interactions between D213 (magenta) and R230 (cyan) and/or Y224 (green). (A,B,D,& E) Time series plots of the D213-Y224/R230 donor-acceptor salt bridge distances of the active and inactive states for both simulated replicates. (C & F) Graphical illustration of salt bridge interactions between the D213-Y224/R230 in the active and inactive states of the CB1 conformation.

## Conclusion

In summary, our comparative MD simulations offer insights into the distinct conformational landscapes of CB1 in active versus inactive states under *apo* conditions. Our multi-microsecond timescale analyses have the potential to enhance our understanding of the conformational mechanisms that facilitate transitions between functional states of this important therapeutic receptor.

A major finding from our study is that contrary to traditional models, inactive CB1 explores a much broader conformational landscape compared to the more confined and rigidified active state ensemble at least under *apo* conditions. The inactive state exhibits substantially increased structural heterogeneity and plasticity in the absence of stabilizing ligand interactions. This was demonstrated by the increasing RMSD values, indicating a continuous deviation from the initial inactive state from the crystal structure. In contrast, the plateauing RMSD profile for the active state reflects its maintenance of a compact conformational state despite ligand unbinding. Our findings reveal the presence of intrinsic malleability unique to the inactive state that may prime CB1 for facile conformational transitions upon ligand binding. We hypothesize the wide structural ensemble adopted by inactive CB1 likely represents intermediate structures that are explored during the activation process.

Importantly, we identified the vital role of TM7 in distinguishing CB1 inactive and active state dynamics. TM7 forms extensive stabilizing intramolecular couplings with surrounding helices including TM2 in the active state that are disrupted when inactive. This enables outward movement and flexibility of TM7 that is uniquely observed in the inactive state simulations. Our findings highlight TM7 as an important helix governing the state-dependent dynamics of CB1 through this specialized switching behavior. The functional significance of TM7 dynamics arises from its known involvement in the activation mechanism of class A GPCRs such as CB1. Rearrangement of TM7 appears to be an early event triggering the switch between functional states. Our findings suggest that the enhanced flexibility of TM7 in the inactive state enables the conformational changes necessary to impede further signaling upon ligand dissociation, thereby promoting stabilization of the inactive state. The simulations provide atomistic resolution into the role of TM7 functional rotation during CB1 activation.

Furthermore, our study revealed extensive networks of electrostatic interactions that form an interconnected system, distinctly characterizing each state. For instance, a dual salt bridge arrangement centered around Lys 192, concurrently pairing with Asp 176 and Asp 184, exhibited remarkable persistence throughout the inactive state simulations but was disrupted in the active state. Our analysis highlights the intricate rearrangement of H-bond and salt bridge networks as integral to the activation process.

Collectively, our findings shed light on the conformational changes driving functional transitions in CB1. By revealing the heightened flexibility of the inactive state, dynamics of TM7, and rearrangement of electrostatic networks, this study elucidates the mechanistic underpinnings of state-dependent modulation of CB1 at an atomistic level. These insights offer valuable context for interpreting novel CB1 crystal structures and may serve as a foundation for developing drugs that selectively target the dynamic conformational changes associated with CB1 function.

## Supporting Information Available

Figures S1–S3 and Movies S1-S3 in Supporting Information provide additional analysis based on our MD simulations as discussed in the manuscript.

## Acknowledgement

The research reported in this work was supported by the National Institute of General Medical Sciences of the National Institutes of Health under award numbers R15GM139140 and R35GM147423, as well as by the National Science Foundation grant CHE 1945465 and the Arkansas Biosciences Institute. This research is part of the Frontera computing project by LRAC at the Texas Advanced Computing Center (TACC) through Grant CHE21003, made possible by the National Science Foundation award OAC-1818253. This work also utilized Stampede by the Extreme Science and Engineering Discovery Environment (allocation MCB150129), which is supported by National Science Foundation grant number ACI-1548562. Additionally, this research was also supported by the Blue Waters sustained-petascale computing project, which is supported by the National Science Foundation (awards OCI-0725070 and ACI-1238993) and the state of Illinois, and by the Arkansas High-Performance Computing Center which is funded through multiple National Science Foundation grants and the Arkansas Economic Development Commission.

