## Supporting Information for "Differential Behavior of Conformational Dynamics in Active and Inactive States of Cannabinoid Receptor 1"

### Supporting Information: Differential Behavior of Conformational Dynamics in Active and Inactive States of Cannabinoid Receptor 1 Revealed by Microsecond-Level Molecular Dynamics Simulation

Ugochi H. Isu, Adithya Polasa, and Mahmoud Moradi*

Department of Chemistry and Biochemistry, University of Arkansas, Fayetteville, AR 72701

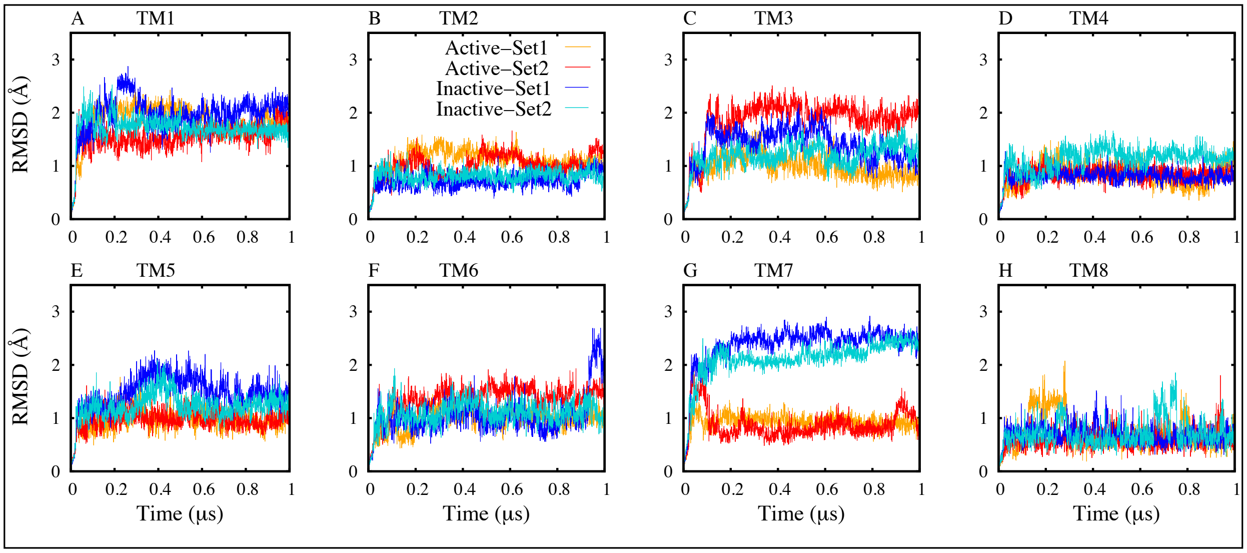


**Fig. S1.** **Cα atom RMSD analysis of TM helices with respect to the initial model across Simulation Sets 1 and 2.**

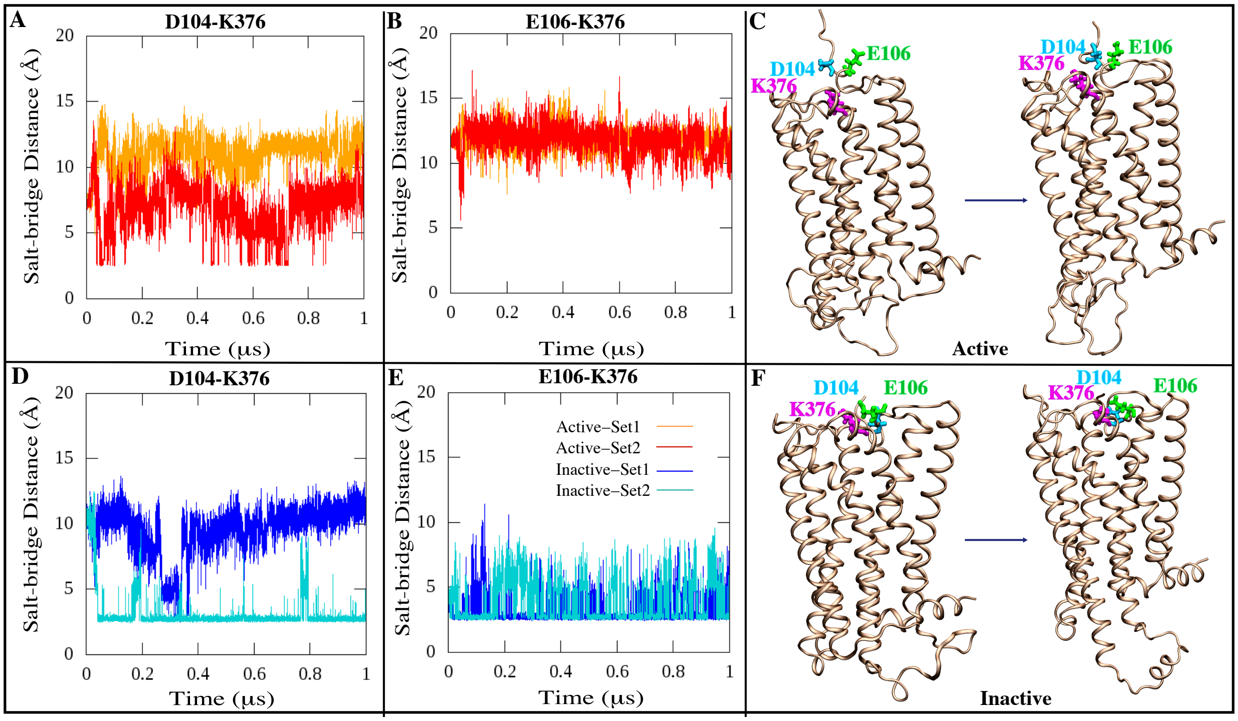


**Fig. S2.** Network of salt bridge interactions partially present in inactive state but absent in active state. Salt bridges between K376 (magenta) and D104 (cyan)/E106 (green). (A, D) Time series plots of the K376-D104, (B, E) Time series plots of the K376-E106, donor-acceptor salt bridge distances of the active and inactive states for both simulated replicates. (C, F) Graphical illustration of salt-bridge interactions between the K376-D104/E106 in the active and inactive states.

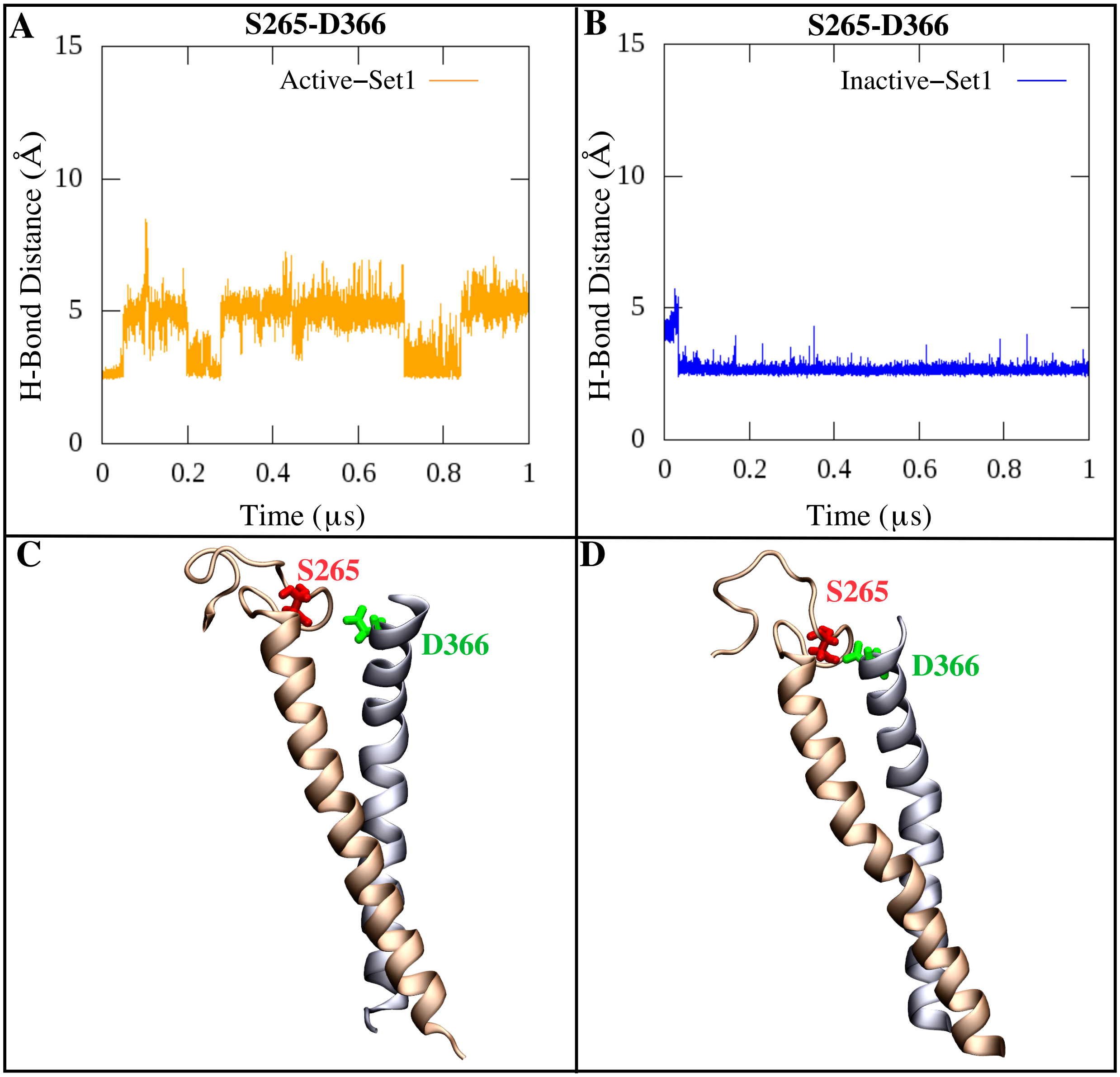


**Fig. S3.** Differential hydrogen bonding between S265 and D366 in (A) active and (B) inactive models of set 1 CB1 simulation.

#### Supporting Movies

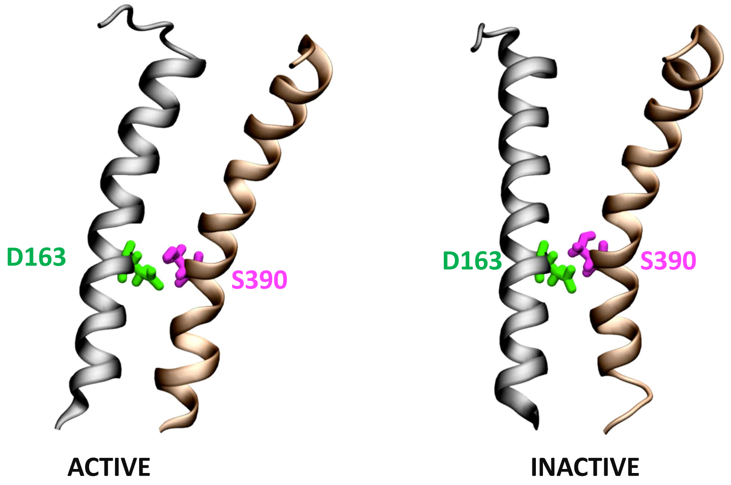


**Mov. S1.** Movie visualizing H-bond interaction between D163 on TM2 and S390 on TM7.

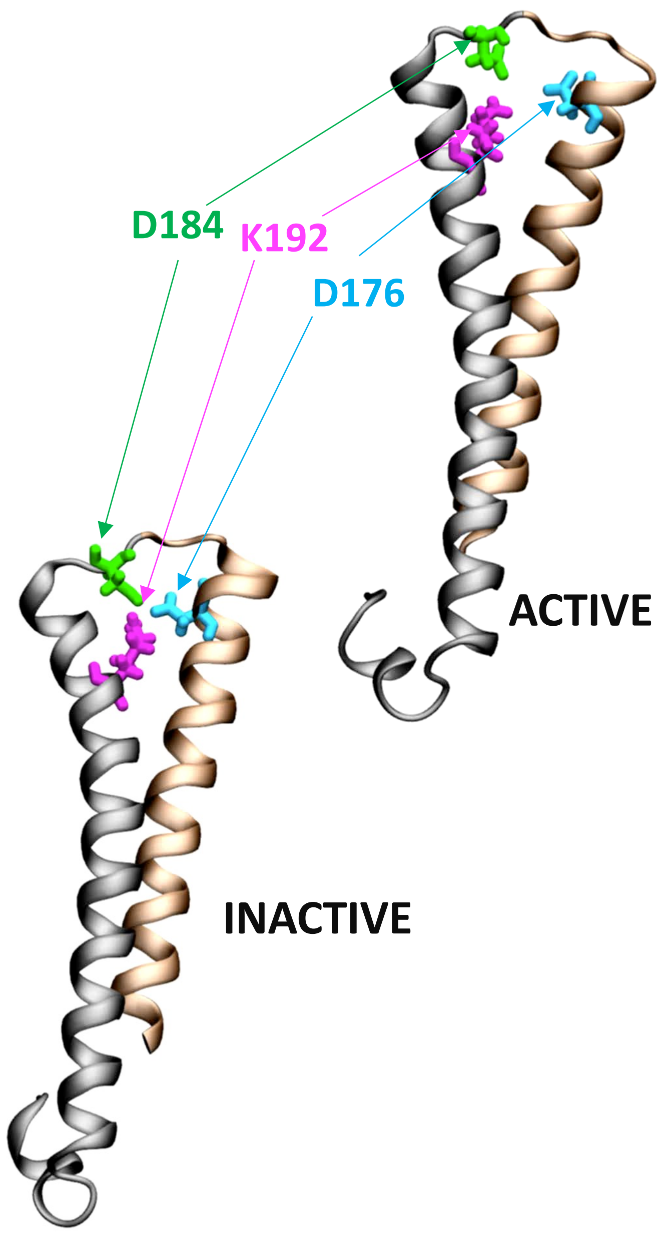


**Mov. S2.** Salt–bridge interaction network between K192 and D176/D184. Positively charged K192 (magenta) fluctuates between negatively charged D176 and D184 (cyan and green) respectively.

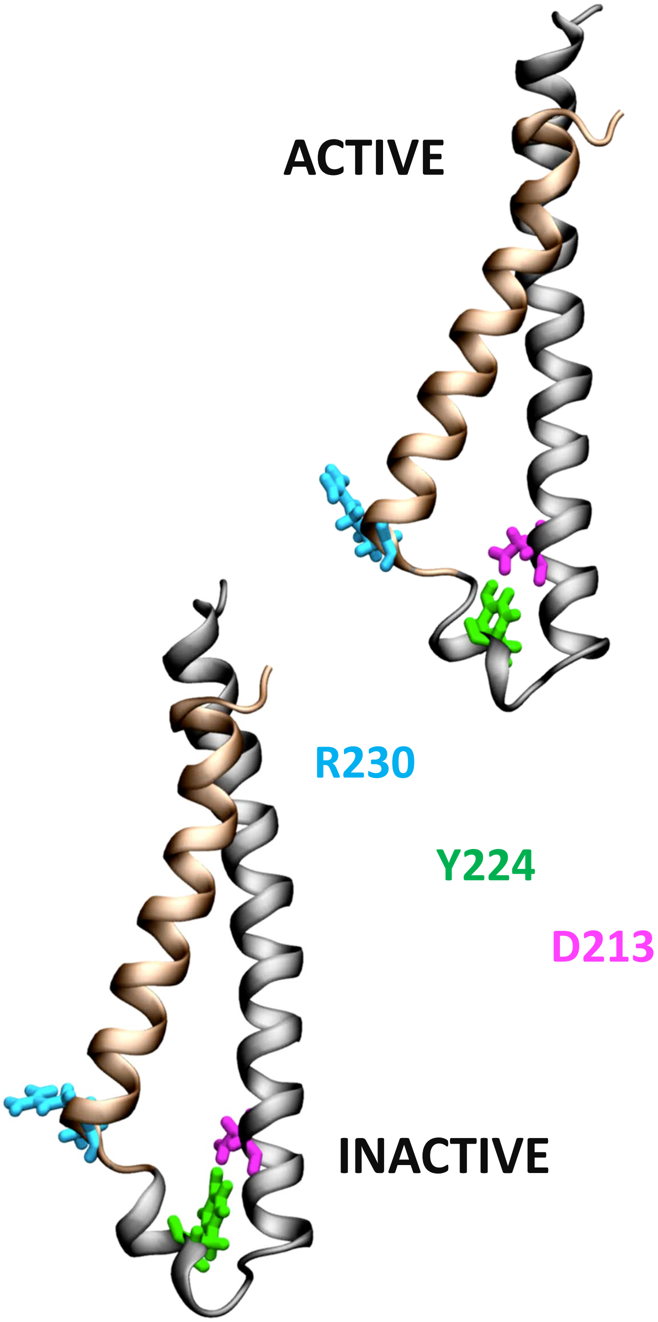


**Mov. S3.** Movie showing interactions between D213, Y224, and R230.
